# A genome-scale metabolic model for *Methylococcus capsulatus* predicts reduced efficiency uphill electron transfer to pMMO

**DOI:** 10.1101/329714

**Authors:** Christian Lieven, Leander A. H. Petersen, Sten Bay Jørgensen, Krist V. Gernaey, Markus J. Herrgard, Nikolaus Sonnenschein

## Abstract

**Background:** Genome-scale metabolic models allow researchers to calculate yields, to predict consumption and production rates, and to study the effect of genetic modifications *in silico*, without running resource-intensive experiments. While these models have become an invaluable tool for optimizing industrial production hosts like *E. coli* and *S. cerevisiae*, few such models exist for one-carbon (C1) metabolizers.

**Results:** Here we present a genome-scale metabolic model for *Methylococcus capsulatus*, a well-studied obligate methanotroph, which has been used as a production strain of single cell protein (SCP). The model was manually curated, and spans a total of 877 metabolites connected via 898 reactions. The inclusion of 730 genes and comprehensive annotations, make this model not only a useful tool for modeling metabolic physiology, but also a centralized knowledge base for *M. capsulatus*. With it, we determined that oxidation of methane by the particulate methane monooxygenase is most likely driven through uphill electron transfer operating at reduced efficiency as this scenario matches best with experimental data from literature.

**Conclusions:** The metabolic model will serve the ongoing fundamental research of C1 metabolism, and pave the way for rational strain design strategies towards improved SCP production processes in *M. capsulatus*.

## Availability of Data and Materials

The metabolic model, scripts and corresponding datasets generated and/or analysed during the current study are available in the GitHub repository.

- Archived: https://github.com/ChristianLieven/memote-m-capsulatus
- Most Recent: https://github.com/ChristianLieven/memote-m-capsulatus/tree/develop

The data that support the biomass equation constructed in this study are available from Unibio at

- http://www.unibio.dk/end-product/chemical-composition-1
- http://www.unibio.dk/end-product/chemical-composition-2

Restrictions may apply to the availability of these data. Data are however available from the authors upon reasonable request and with permission of Unibio.

## Background

The Gram-negative, obligate-aerobe *Methylococcus capsulatus* is a methane oxidizing, gamma-proteobacterium. Since its initial isolation by Foster and Davis [1], the organism has been subject to a wide array of studies. The global role of *M. capsulatus* as a participant in the carbon cycle has been elucidated [2, 3] as well as its effects on human [4] and animal health and disease [5]. Specifically the latter studies have been triggered by a considerable commercial interest in *M. capsulatus* as the primary microbe used for the production of Single Cell Protein (SCP) as animal feed starting in the 70s [6]. Now that hydraulic fracturing has made natural gas a cheap and abundant feedstock [7], the application of *M. capsulatus* for this purpose is being explored again [8, 9]. Another portion of studies, however, has focused on uncovering the biochemical and genetic basis of the organism’s unique metabolism [10]. Yet, the greatest interest has been the role and function of the initial enzyme in methanotrophy, methane monooxygenase [11], responsible for oxidation of methane to methanol.

*Methylococcus capsulatus* is able to express two distinct types of methane monooxygenases: a soluble form of methane monooxygenase (sMMO) and a particulate, or membrane-bound form (pMMO). The expression of these enzymes is strongly influenced by the extracellular concentration of copper; when *M. capsulatus* is grown in the presence of low levels of copper the sMMO is active, while the pMMO is predominantly active at high levels. Both enzymes require an external electron donor to break a C-H bond in methane. While the electron donor for the sMMO is NADH [12–15], the native reductant to the pMMO has not yet been elucidated due to difficulties to purify the enzyme and assay its activity *in vitro* [11]. Three hypotheses regarding the mode of electron transfer to the pMMO have been suggested previously: 1) formaldehyde oxidation provides the electrons required to oxidize methane in the form of NADH, while the electrons that are released from the oxidation of methanol are passed via a cytochrome ‘redox arm’ to a terminal oxidase, generating ATP. 2) A direct coupling between the pMMO and the methanol dehydrogenase (MDH) allows an efficient exchange of electrons, leading to the latter reaction driving the former. 3) Finally, it is possible that electrons are transferred to the pMMO from MDH through the ubiquinol pool by the action of a reversely operating ubiquinol-cytochrome-c reductase. This mode is referred to as ‘uphill’ electron transfer [17].

A genome-scale metabolic model (GEM) not only represents a knowledge base that combines the available physiological, genetic and biochemical information of a single organism [16], but also provides a testbed for rapid prototyping of a given hypothesis [17]. Hence, we present the first manually curated genome-scale metabolic model for *Methylococcus capsulatus,* with the intent of supplying the basis for hypothesis-driven metabolic discovery and clearing the way for future efforts in metabolic engineering [18]. Using the GEM, we investigate the nature of electron transfer in *M*. *capsulatus* by comparing the model’s predictions against experimental data from Leak and Dalton [19]. Furthermore, we compare its predictions to those of the model of *M. buryatense* [20] and explain notable differences by considering the proposed electron transfer modes.

## Results and Discussion

### Reconstruction

The presented genome-scale metabolic reconstruction of *M. capsulatus* termed iMcBath was based on BMID000000141026, an automatic draft published as part of the Path2Models project [21]. The whole genome sequence of *Methylococcus capsulatus* (Bath) (GenBank AE017282.2 [22]) was used to aid the curation process and to supply annotations (see material & methods). This metabolic reconstruction consists of 840 enzymatic reactions that interconvert 783 unique metabolites. The total number of reactions, including exchange, demand and the biomass reactions, is 892. The model attributes reactions and metabolites to three distinct compartments: The cytosol, the periplasm, and the medium that is referred to as extracellular in the model. Gene-Protein-Reaction rules (GPR) associated with 730 unique genes support 85.5% of the included reactions with, leaving 122 reactions without a GPR. The GPRs include representation of 24 enzyme complexes.

The model is made available in the community-standard SMBL format (Level 3 Version 2 with FBC).

### Biomass reaction

In stoichiometric models biological growth is represented as the flux through a special demand reaction. The so-called biomass reaction functions as a drain of metabolites, which are either highly reduced non-polymeric macromolecules such as lipids and steroids or precursors to typical biopolymers such as nucleic acids, proteins or long-chain carbohydrates. The stoichiometry of an individual precursor was calculated from the principal composition of *M. capsulatus* as reported by Unibio [23]. The monomer composition of individual macromolecules was calculated from different sources. A detailed account of the resources is provided in the methods section and an overview of the biomass reaction is given in Supplement Table 1.

**Table 1.**
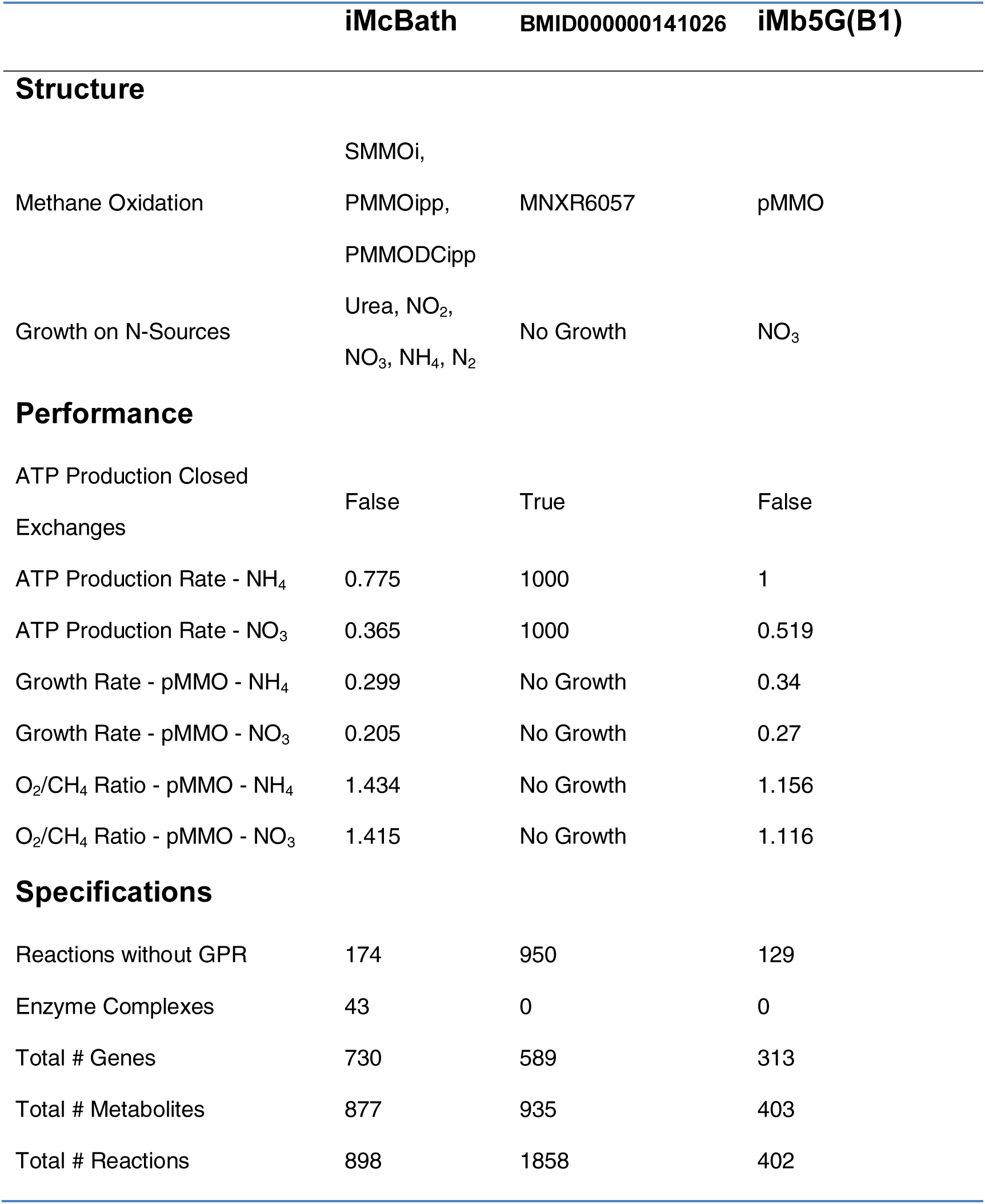
Model Comparison. The presented reconstruction iMcBath, the automated draft BMID000000141026 and iMb5G(B1), a genome-scale reconstruction of *Methylomicrobium buryatense* strain 5G(B1).

The growth-associated maintenance (GAM) and the non-growth associated maintenance (NGAM) requirements for *M. capsulatus* are yet to be determined experimentally. Therefore, a GAM value of 23.087 mmol ATP gDW^-1^ h^-1^ was estimated according to Thiele et al. [16] based on the data for *E. coli* published by Neidhardt et al. [24]. The value for GAM is expected to increase with the growth rate of the cells [25]. Like de la Torre et al. had done for *M. buryatense* [20], we assumed the non-growth associated maintenance (NGAM) of *M. capsulatus* to be similar to that of *E. coli* thus setting it to 8.39 mmol ATP gDW^-1^ h^-1^ [26].

### Metabolism

Much of the focus in curating the initial draft model BMID000000141026 was put on the central carbon metabolism and respiration of *M. capsulatus*.

Three possible modes of electron transfer to the pMMO have been proposed.

1) In the *redox-arm* mode [27], the methanol dehydrogenase (MDH) passes electrons via cytochrome c555 (cL) [28] and cytochrome c553 (cH) [29] to either a CBD-or AA3-type terminal oxidase [30] and thus contributes to building up a proton motive force (PMF) and the synthesis of ATP (Figure 1). The electrons required for the oxidation of methane are provided through ubiquinone (Q8H2), which in turn received electrons from various dehydrogenases involved in the oxidation of formaldehyde to CO_2_. Although no binding site has been identified, there is support for pMMO reduction by endogenous quinols [31].

**Figure 1.**
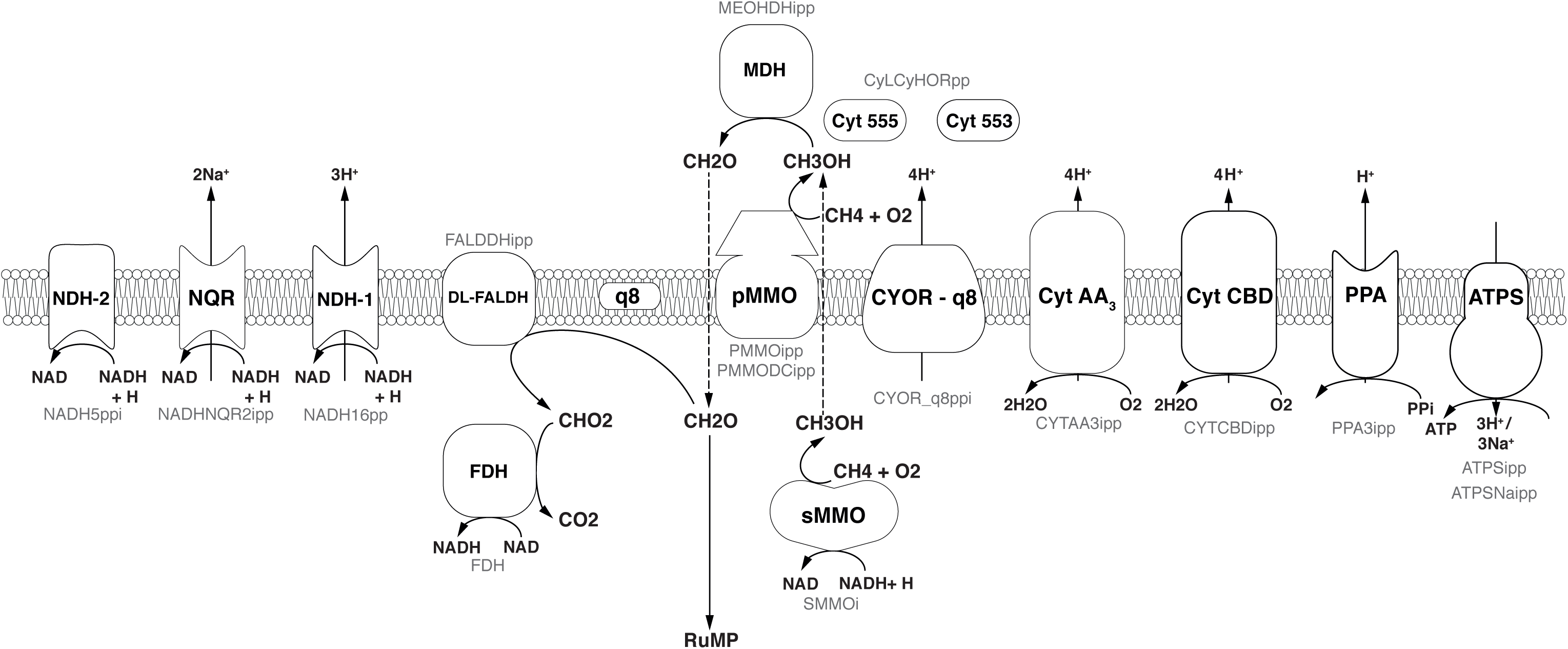
Overview of the respiratory chain in *Methylococcus capsulatus* as implemented in iMcBath. Black text in the center of the symbols denotes the common abbreviation, while the faint grey text below denotes the corresponding reaction ID in the metabolic model.

2) With the *direct coupling* mode, the MDH is able to directly pass electrons to the pMMO [32–34]. Here, cytochrome c555 is the electron donor to the pMMO instead of ubiquinol. This mode is supported by results from cryoelectron microscopy which indicates that the pMMO and the MDH form a structural enzyme complex [35].

3) The *uphill electron transfer* mode supposes that the electrons from cytochrome c553 can reach the ubiquinol-pool facilitated by the PMF at the level of the ubiquinol-cytochrome-c reductase. This mode was proposed by Leak and Dalton [36] as it could explain the observed reduced efficiency.

To determine which of these modes is most likely active in *M. capsulatus,* Leak and Dalton [36] developed mathematical models for each mode based on previous calculations [37, 38]. However, their models did not reflect the experimental results. De la Torre et al. constructed a genome-scale metabolic model (GEM) for *Methylomicrobium buryatense* to investigate growth yields and energy requirements in different conditions [20]. They found that the *redox-arm* mode correlated least with their experimental data for *M. buryatense*.

To study which of the three modes of electron transfer is active in *M. capsulatus*, they were each implemented in the model. The implementation is illustrated in Figure 2. To include the *redox-arm* we implemented the reaction representing the particulate methane monooxygenase, in the model termed as PMMOipp, utilizing Q8H2 as a cofactor. Accordingly, a variant of the pMMO reaction was added to account for a possible direct coupling to the MDH. In this variant reaction, termed PMMODCipp, the cofactor is cytochrome c555, represented as the metabolite focytcc555_p in the model. To enable an *uphill electron transfer*, the reaction representing the ubiquinol-cytochrome-c reductase (CYOR_q8ppi in the model), was constrained to be reversible while keeping PMMOipp active.

**Figure 2.**
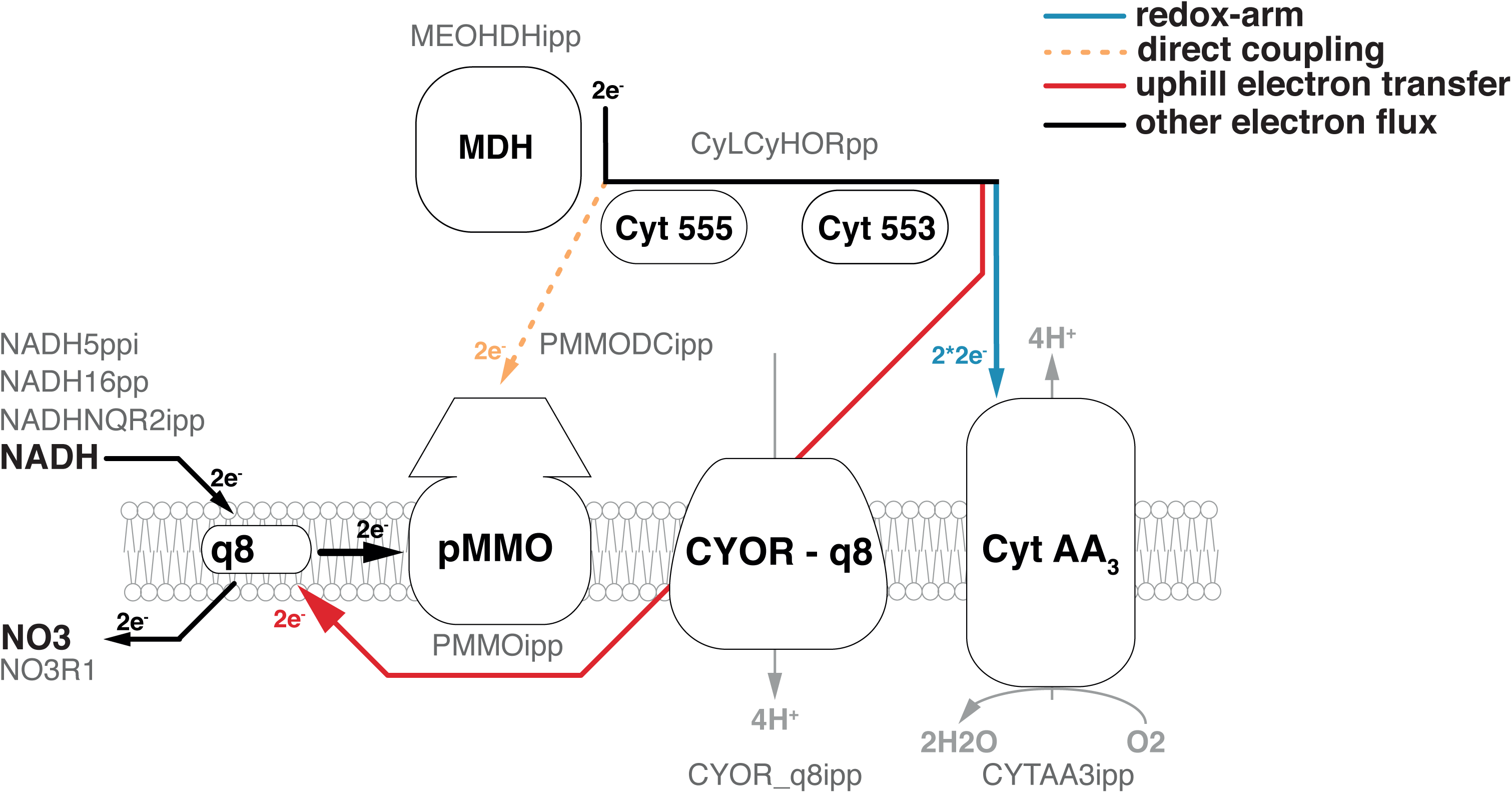
The three possible modes of electron transfer to the pMMO mapped onto an excerpt of the respiratory chain. 1) Redox-arm: The methanol dehydrogenase transfers electrons to the terminal oxidase, while the pMMO draws electrons from the quinone pool. 2) Direct coupling: Electrons from the oxidation of methanol are transferred directly to the pMMO. 3) Uphill electron transfer: Electrons from the methanol dehydrogenase feed back into the ubiquinol-pool. Black text denotes the common name, while faint grey text denotes the reaction ID in the metabolic model.

Following the path of carbon through metabolism downstream from the MDH, the model includes both the reaction for a ubiquinone-dependent formaldehyde dehydrogenase [39], termed FALDDHipp, and an NAD-dependent version, termed ALDD1 (Figure 3). Despite of the initial evidence for the latter reaction [40] having been dispelled by Adeosun et al. [41], it was added to allow further investigation into the presence of a putative enzyme of that function. An additional pathway for formaldehyde oxidation represented in both genome and model is the tetrahydromethanopterin(THMPT)-linked pathway [42].

**Figure 3.**
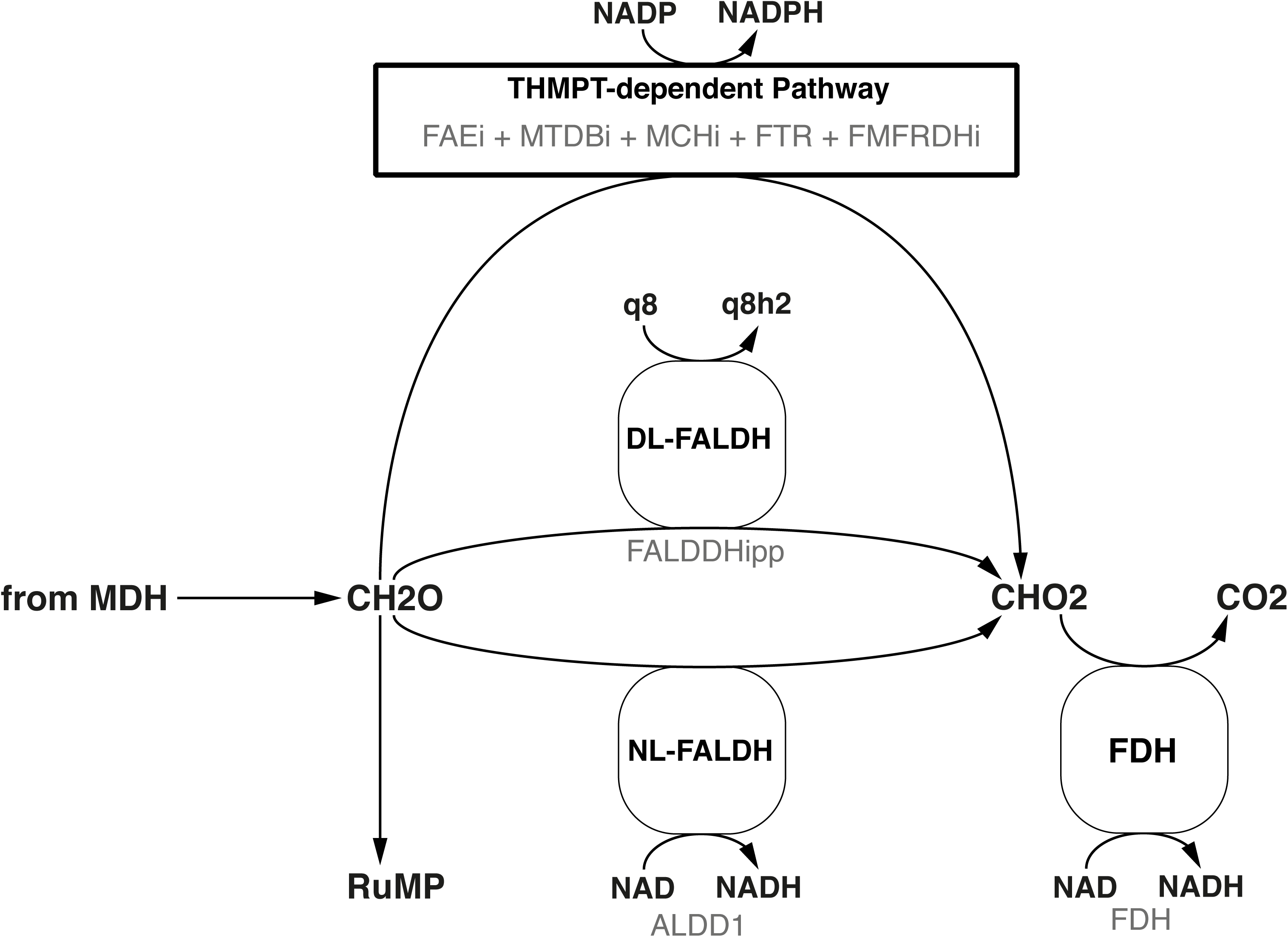
Three different formaldehyde oxidation pathways are represented in the model. Black text denotes the common name, while faint grey text denotes the reaction ID in the metabolic model.

Formaldehyde assimilation in *M. capsulatus* occurs primarily through the ribulose monophosphate (RuMP)-pathway. As outlined by Anthony [10], the RuMP-pathway has four hypothetical variants. Based on the annotated genome published by Ward et al. [22], we identified not only both C6 cleavage pathways depending either on the 2-keto, 3-deoxy, 6-phosphogluconate (KDPG) aldolase (EDA) or the fructose bisphosphate aldolase (FBA), but also the transaldolase (TA) involved in the rearrangement phase that regenerates ribulose 5-phosphate. The alternative to a transaldolase-driven rearrangement step is the use of a sedoheptulose bisphosphatase, which was not included in the initial annotation. Strøm et al. could not detect specific activity using cell-free preparations [43]. Yet, we decided to add a corresponding reaction for two reasons. First, the FBA has been characterized to reversibly catalyze sedoheptulose bisphosphate cleavage [44], which is reflected by the reaction FBA3 in the model. Second, the pyrophosphate-dependent 6-phosphofructokinase (PFK_3_ppi) was reported to have higher affinity and activity to the reversible phosphorylation of seduheptulose phosphate than compared to fructose 6-phosphate [45]. Thus, all of the resulting four combinations that make up the RuMP-pathway are represented in this metabolic model (Figure 4).

**Figure 4.**
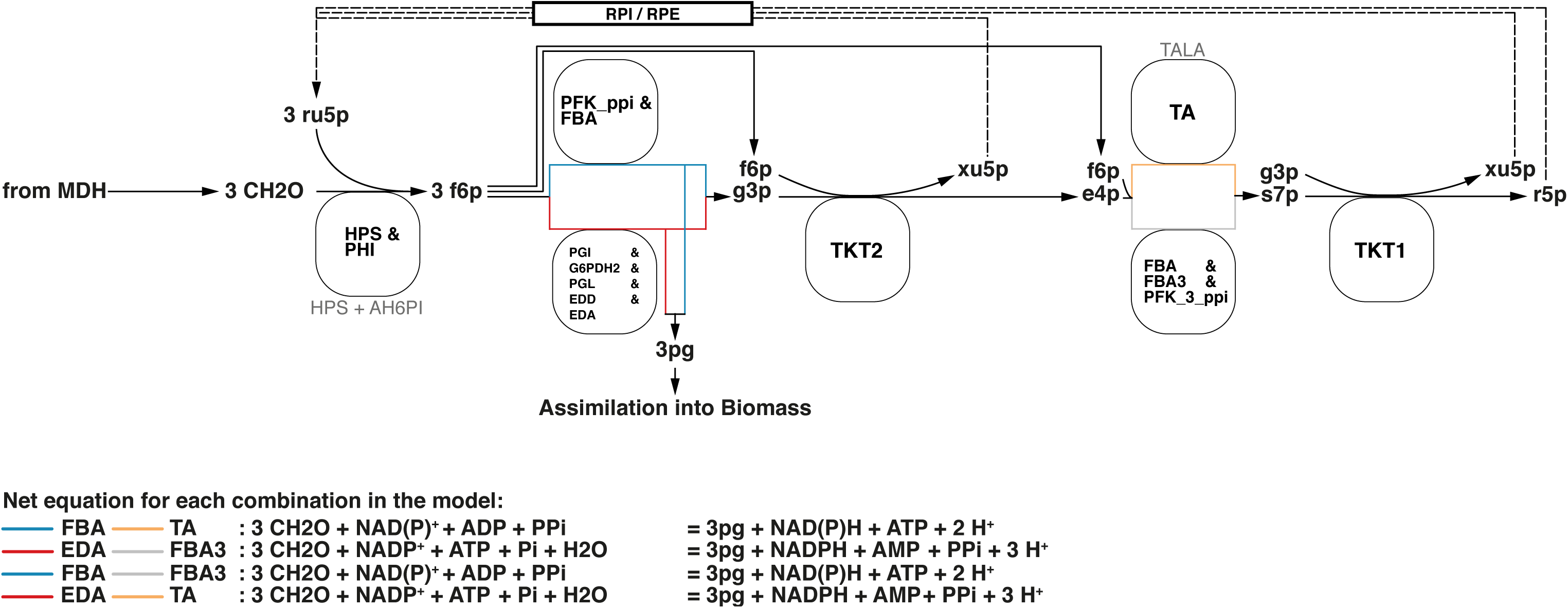
All four variants of the RuMP pathway are represented in the metabolic model. If there is a common enzyme name the black text denotes the common name, while faint grey text denotes the reaction ID in the metabolic model. Otherwise the black text is also the reaction ID in the model.

The genome of *Methylococcus capsulatus* encodes genes for a complete Calvin Benson Bassham (CBB) cycle [46–48] and a partial Serine pathway for formaldehyde assimilation [22]. It was argued by Taylor et al. [49] that *M. capsulatus can metabolize* glycolate (a product of the oxygenase activity of the ribulose bisphosphate carboxylase (RBPC)) via this pathway. Both Taylor and Ward, further suggested the presence of unique key enzymes to complete the Serine pathway, such as hydroxymethyl transferase, hydroxypyruvate reductase and malate-CoA lyase. However, since the gene annotation did not reflect this and the RuMP pathway is reportedly the main pathway for formaldehyde assimilation [50], these putative reactions were not included.

All genes of the TCA cycle were identified in the genome sequence and all associated reactions were included accordingly [22]. Because no activity of the 2-oxoglutarate dehydrogenase has so far been measured *in vivo* [50, 51], the associated reactions have been constrained to be blocked (both lower and upper bounds were set to zero). This way they can easily be activated if needed. For instance, if a growth condition is discovered where activity for this enzyme can be detected.

Based on reactions already present in BMID000000141026, the information in the genome annotation, and the measured biomass composition, we curated the biosynthetic pathways of all proteinogenic amino acids, nucleotides, fatty acids, phospholipids, panthothenate, coenzyme A, NAD, FAD, FMN, riboflavin, thiamine, myo-inositol, heme, folate, cobalamine, glutathione, squalene, lanosterol, peptidoglycan. Since no corresponding genes could be identified, reactions catalyzing the biosynthesis of lipopolysaccharide (LPS) were adopted from iJO1366 [52] under the assumption that the biosynthesis steps among gram-negative bacteria require amounts of ATP comparable to *Escherichia coli*.

*Methylococcus capsulatus* is able to metabolize the nitrogen sources ammonium (NH_4_) and nitrate (NO_3_) in a variety of ways. When the extracellular concentration of NH_4_ is high, alanine dehydrogenase (ADH) is the primary pathway for nitrogen assimilation into biomass [53]. In addition, the two monooxygenases are able to oxidize ammonium to hydroxylamine [15, 54], which is then further oxidized by specific enzymes first to nitrite [55] (Figure 5), and even to dinitrogen oxide [56]. NO_3_ is reduced to NH_4_ via nitrite and ultimately assimilated via the glutamine synthetase/ glutamine synthase (GS/GOGAT) pathway. Furthermore, it has been shown that *M. capsulatus* is able to fix atmospheric nitrogen (N_2_) [57]. The nitrogenase gene cluster has been identified [58] and annotated accordingly [22], and the corresponding reactions have been included in the model. As the enzyme has not yet been specifically characterized, the nitrogenase reaction was adapted from iAF987 [59]. A schema showing these reactions side-by-side is displayed in Figure 6.

**Figure 5.**
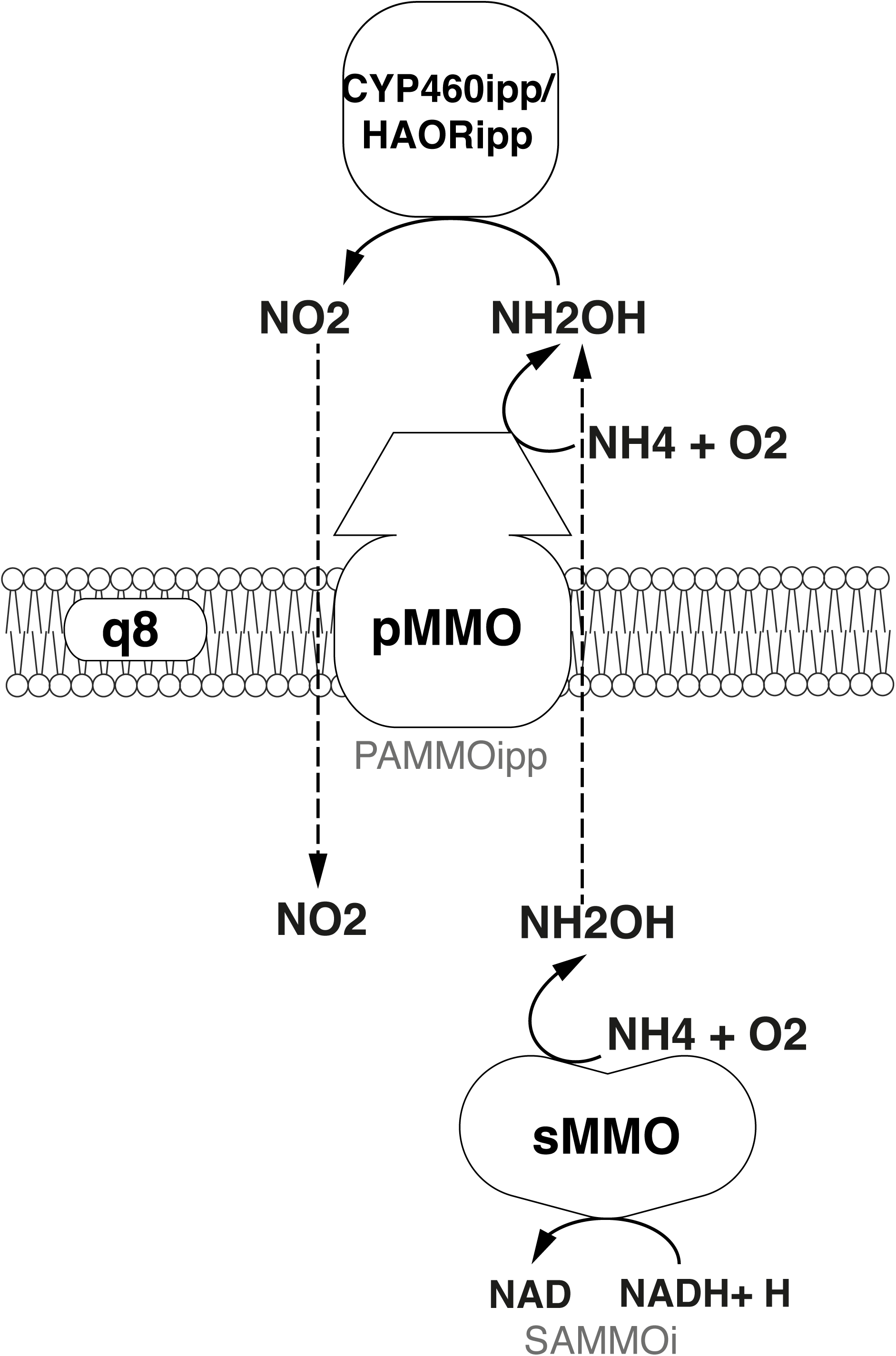
Oxidation of ammonia to nitrite.

**Figure 6.**
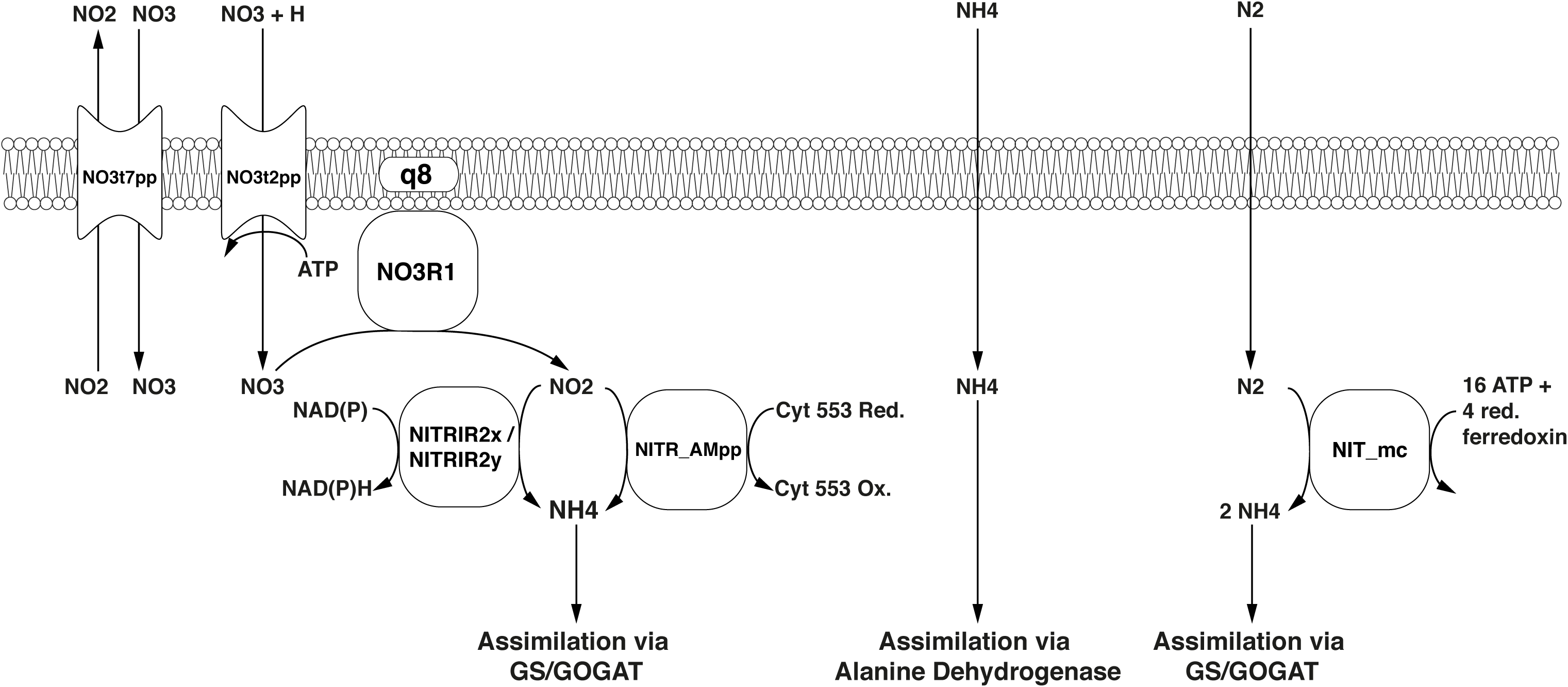
Nitrogen assimilation and fixation pathways represented in the model. Black text in the centre of the symbols denotes reaction IDs. Common names are used for metabolites.

A metabolic map of all the reactions in the model was built using Escher [60] and is available as Supplement Figure 1.

### Extension and manual curation

Starting reconstruction on the basis of an automated draft required additional effort to be able to use it for calculations. The automated draft used two sets of ID namespaces, BiGG [61] and MetaNetX (MNX) [62]. Hence, the first step in the curation efforts consisted of unifying the namespace by mapping all metabolite and reaction identifiers from MNX into the human-readable BiGG namespace. Individual metabolites and reactions with unmappable IDs, that could not be identified in the BiGG database and for which there was little evidence in the form of GPR rules, were removed from the model. Several metabolite formulas contained polymer units, and many reactions lacked EC numbers. Using the MNX spreadsheet exports ‘chem_prop.tsv’ and ‘reac_prop.tsv’ from version 1.0 [63] and 2.0 [64] most of these issues were resolved. Due to said malformed and missing metabolite formulae, many reactions were not mass-charge-balanced. We used the ‘check_mass_balance’ function in cobrapy [65] to identify and balance 99.4% of the reactions in the model. The remaining five reactions, belonging to fatty acid biosynthesis, involve a lumped metabolite that represents the average fatty acid content of *M. capsulatus*.

Another challenge with extending the automated draft was that many reactions were inverted meaning that reactants and products were switched, and constrained to be irreversible. Consulting the corresponding reactions in MetaCyc [66], these instances were corrected manually. For each precursor in the biomass reaction, the corresponding biosynthetic pathway was gap-filled and manually curated. To identify the appropriate pathways, MetaCyc pathway maps from the *M. capsulatus* Bath specific database were used to compare the reactions that were present in the draft reconstruction (https://biocyc.org/organism-summary?object=MCAP243233).

In order to enable other researchers to integrate iMcBath into their respective workflows and to simplify model cross-comparisons, we included MIRIAM-compliant annotations for a majority of the model’s reactions (96%) and metabolites (98%) [67].

As a last step we added transport reactions that were not in the draft reconstruction. Inferring membrane transport reactions from the genome sequence is difficult, as usually the precise 3D structure of transport proteins dictates which metabolite classes can be transported [68]. Even if the substrates are known, the energy requirements of transport are often undefined. Working on protein sequence matches using PsortB 3.0 [69], combined with BLAST [70] matches against TransportDB [71] and the Transporter Classification Database (TCDB) [72], we were able to identify 56 additional transport reactions. We have limited the number of transporters to be added, focusing specifically on transporters with known mechanisms and transport of metabolites already included in the model. A list of putative transport-associated genes that we did not consider is available at https://github.com/ChristianLieven/memote-m-capsulatus. Some of these genes were reported by Ward et al. to facilitate the uptake of sugars [22]. Kelly et al. suggest [50] these genes may allow a function similar to *Nitrosomas europaea,* which is able grow on fructose as a carbon-source with energy from ammonia oxidation [73]. This list is a good starting point to study potential alternate carbon source use by *M. capsulatus*.

### Validation of the Model

To determine which combination of the three aforementioned electron transfer modes is active in *Methylococcus capsulatus*, we constrained the model based on experiments conducted by Leak and Dalton [19]. Since the three modes relate to how the pMMO receives electrons, we focused on the data generated by growing *M. capsulatus* in high-copper medium, which is the condition in which pMMO is predominantly active. We used the average of carbon and oxygen-limited measurements as a reference. Having constrained the model, we compared the Leak and Dalton’s measurements for the ratio of oxygen consumption to methane consumption (O_2_/CH_4_) to the predictions of the model (Figure 7A). We considered the O_2_/CH_4_ ratio to be a key metric for the respiratory chain in *M. capsulatus*, as it is a function of the mode of electron transfer to the pMMO. The central carbon metabolism was left unconstrained.

**Figure 7.**
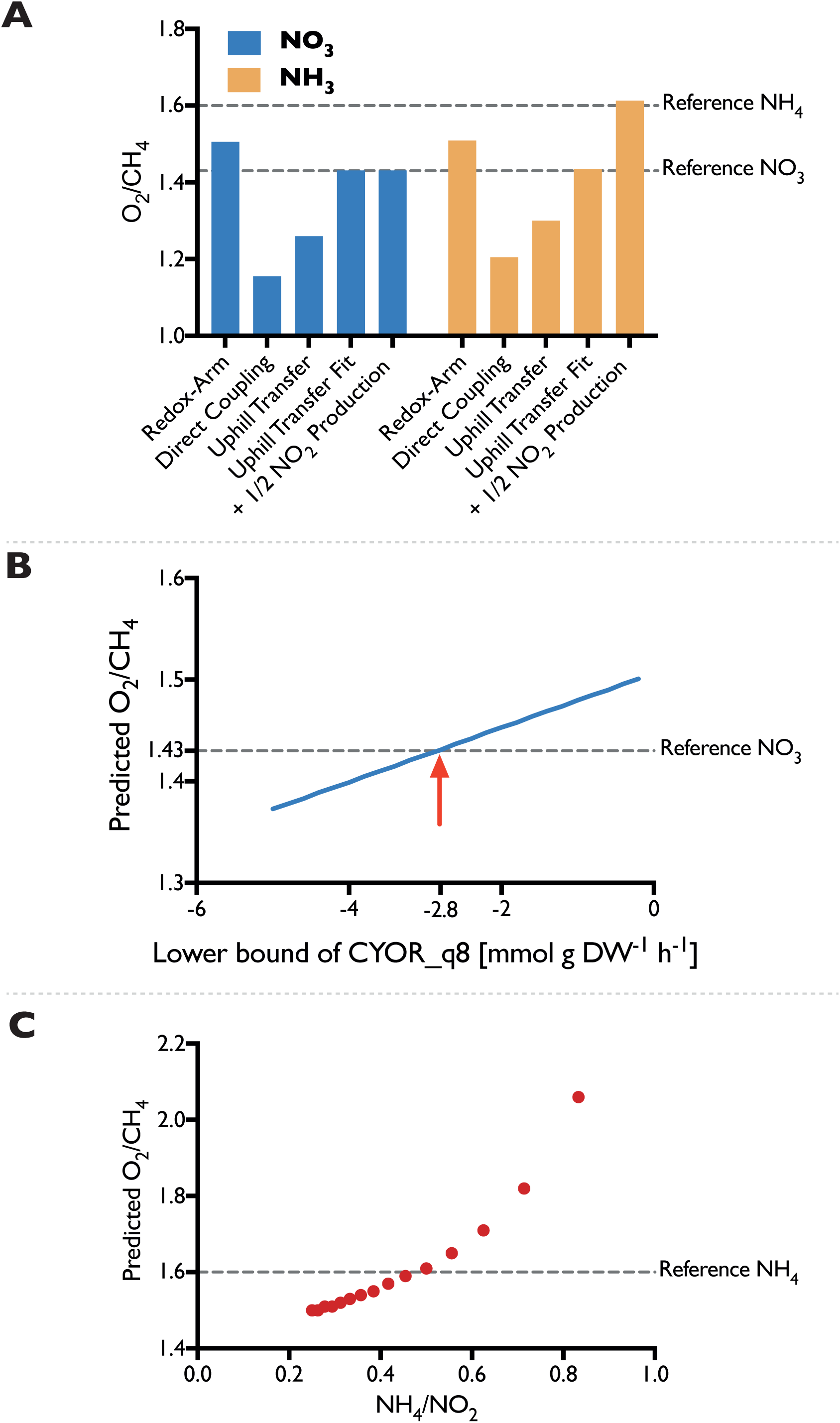
Validation of iMcBath. A: Using each of the three electron transfer modes exclusively (Redox-Arm, Direct Coupling, Uphill Transfer), the ratios of oxygen to methane consumption (O_2_/CH_4_) predicted by iMcBath were compared to experimental values from Leak and Dalton [19]. Since none of the three modes matched the reference, the efficiency of the Uphill Transfer was reduced by iteratively constraining the flux through the ubiquinol-cytochrome c reductase (CYOR_q8) reaction as shown in **B.** Using a lower bound of −2.8 improved the fit to the reference value for cells grown on NO_3_ (**A** – Uphill Transfer Fit) **C:** To account for the energy loss through NH_3_ oxidation, several ratios of NH_4_ uptake to NO_2_ production were considered. The closest fit was achieved with a ratio of around 0.5 (**A** – + ½ NO_2_ Production)

Under the assumption that the mode of electron transfer to pMMO would be independent of the source of nitrogen, we compared the O_2_/CH_4_ ratios of the model constrained to employ one of the three modes of electron transfer exclusively to the corresponding reference values of *M. capsulatus* grown on either NO_3_ or NH_4_. However, neither of the modes adequately represented the measured O_2_/CH_4_ ratios of about 1.43 and 1.6, respectively. Although Leak and Dalton had proposed that the *reverse* or *uphill electron transfer* is the most probable mode [36], the model predictions allowing for an unbounded *uphill transfer* did not support this (Figure 7), as the efficiency was almost comparable to predictions using *direct coupling*.

We altered the efficiency of the three modes to determine whether the fit could be improved. For the *redox arm*, we gradually decreased the mol protons required for the synthesis of 1 mol ATP, thereby improving the efficiency. This did not change the O_2_/CH_4_ ratio (See Supplement Table 2). We decreased the efficiency of the *direct coupling* mode (PMMODCipp) by forcing a portion of the flux through the regular pMMO reaction (PMMOipp) using ratio constraints. Although this affected the O_2_/CH_4_ ratio, to achieve a ratio representing the measured value of 1.43 would require a large decrease in efficiency with almost 85% of the incoming carbon to be routed through the regular pMMO reaction (See Supplement Table 2). Because of this, we consider a *direct coupling* to be possible, albeit unlikely. Lastly, we iteratively constrained the lower bound of the reaction associated with the ubiquinol-cytochrome-c reductase (CYOR_q8ppi), to reduce the efficiency of the *uphill-electron transfer*.

As Figure 7B shows, by constraining the reverse flux through this reaction, it is possible to modulate the ratio of O_2_/CH_4_ consumption. We decided to fit the lower bound of ubiquinol-cytochrome-c reductase only to the reference O_2_/CH_4_ ratio of 1.43 with cells grown on NO_3_, in order to avoid an overlap with the effects of NO_2_ production. Leak and Dalton pointed out, that the unexpectedly high O_2_/CH_4_ ratio of 1.6 was the product of latent NH_4_ oxidation rather than assimilation [19] leading to elevated levels of NO_2_. They were uncertain whether this increase could be attributed to the energetic burden of reducing NH_4_ or the uncoupling effect of NO_2_.

To investigate this effect, we introduced a ratio constraint (see Methods) coupling the uptake of NH_4_ to the excretion of NO_2_ and explored a number of values for this ratio (Figure 7C). According to the simulations, the energy spent on reducing about 50% of incoming NH_4_ to NO_2_ is sufficient to account for the observed, high O_2_/CH_4_ ratio of 1.6. Although this shows that the loss of energy could be significant enough to account for the increased ratio, this does not exclude a potential combined effect because of energy decoupling. Regardless it shows that predictions using the metabolic model can accurately reflect the *in vivo* behavior of *M. capsulatus*.

### Comparison with other models

We compared iMcBath, the preceding automated draft reconstruction BMID000000141026, and a genome-scale metabolic model of the gram-negative, gamma-proteobacterium *Methylomicrobium buryatense* strain 5G(B1) [20] to illustrate how much the model has progressed from the draft through manual curation and how the mode of electron transfer affects growth parameters within the group of gamma-proteobacteria (See Table 1).

Unsurprisingly, the automated draft generally performs quite poorly in comparison with the curated models. It’s not possible to produce biomass from methane, even in rich media conditions, which is indicative of gaps in the pathways leading towards individual biomass components. Moreover, ATP can be produced without the consumption of substrate, which in turn means that key energy reactions are not constrained thermodynamically to have the correct reversibility. In the draft, 51% of the reactions are not supported by Gene-Protein-Reaction rules (GPR), while in iMb5G(B1) this percentage is only 32% and in iMcBath the percentage of these reactions is under 20%. GPRs allow modelers to distinguish between isozymes and enzyme complexes by connecting gene IDs either through boolean ‘OR’ or ‘AND’ rules. In iMcBath, 25 complexes in total were curated and formulated as GPR. Neither the draft model nor iMb5G(B1) make this distinction.

In the automated draft, the oxidation of methane was only possible through a reaction representing the sMMO (MNXR6057). In iMcBath, this was corrected by also implementing reactions that represent the pMMO, one with ubiquinone as the electron donor (PMMOipp), and another that receives electrons from the MDH (PMMODCipp). *Methylococcus capsulatus*, like *Methylomicrobium buryatense*, expresses both the soluble and the particulate monooxygenase depending on the availability of copper in the medium. In iMb5G(B1) however only the particulate monooxygenase is represented by the reaction “pMMO”. The ability of *M. capsulatus* to grow on ammonia, nitrate and nitrogen has been characterized experimentally [53]. In addition to that, however, iMcBath also predicts growth on nitrite and urea. Nitrite is an intermediate in the assimilation of nitrate, yet also reported to elicit toxic effects [19, 74], hence *in vivo* growth may only be possible on low concentrations. Growth on urea is possible in the model since we identified *MCA1662* as an ATP-driven urea transporter and gap-filled a urease reaction when curating the cobalamine biosynthesis pathway. As a consequence of this, urea can be taken up into the cytosol and broken down into ammonia and carbon dioxide, the former of which is then assimilated. Further experimental work is necessary to verify this *in vivo*. *M. buryatense* 5G(B1) is reported to grow on nitrate, ammonia and urea, yet without adding the respective transport reactions iMb5G(B1) only grows on nitrate. While the draft model for *M. capsulatus* contains exchange reactions for all the tested nitrogen sources except for urea, it couldn’t grow at all, which again is likely because of gaps in the pathways leading to biomass precursor metabolites.

The difference between *M. capsulatus* and *M. buryatense* becomes apparent when comparing the growth energetics of iMcBath and iMb5G(B1). iMb5G(B1) produces more mmol gDW^-1^ h^-1^ ATP than iMcBath. Because of this the hypothetical growth-rates predicted by iMb5G(B1) are higher than those of iMcBath regardless of the respective nitrogen source. This difference is likely a direct consequence of the mode of electron transfer to the monooxygenase and thus the efficiency of the methane oxidation reaction in total. When comparing the ratio of the uptake rate of oxygen and the uptake rate of methane for the two models, we can see that the resulting values in iMb5G(B1) are lower than in iMcBath. It was recently reported, that instead of the reverse-electron transfer and redox-arm mode active in *M. capsulatus,* a mixture of *direct coupling* from pMMO to MDH and *reverse electron transfer* seems to be the most likely mode in *M. buryatense* 5G(B1) [20].

## Conclusion

IMCBATH is the first, manually curated, genome-scale metabolic model for *Methylococcus capsulatus.* With iMcBath, we combine biochemical knowledge of half a century of research on *Methylococcus capsulatus* into a single powerful resource, providing the basis for targeted metabolic engineering, process design and optimization, omic-data contextualization and comparative analyses. We applied the metabolic model to study the complex electron transfer chain of *M. capsulatus*, by analyzing the three modes of electron transfer that had been proposed previously [36]. We did so by corresponding each mode with the flux through a reaction in the model, and consequently comparing the predicted O_2_/CH_4_ ratios to experimentally measured values by Leak and Dalton [19]. Simulation and experiment agreed either when the model was constrained to employ the *uphill electron transfer* at reduced efficiency or *direct coupling* at strongly reduced efficiency for *M. capsulatus* grown *in silico* on NO_3_ as the source of nitrogen. Although an *uphill electron-transfer* seems more likely, further experimental validation is required as neither mode can be ruled out exclusively. The experimentally observed effect of NH_4_ oxidation to NO_2_ could be replicated by considering the energy burden alone.

Future applications of the metabolic model could include hypothesis testing of the regulation of the MMO in other growth conditions [50], studying the effects of the predicted hydrogenases on the energy metabolism of *M. capsulatus* [22], and exploring venues of metabolic engineering for an improved production of metabolites [75]. Another

## Methods

### Model Curation

After mapping the reaction and metabolite identifiers from the MetaNetX namespace to the BiGG namespace, we proceeded with the curation efforts as follows: First, we chose a subsystem of interest, then we picked a pathway and using information from either the genome sequence, published articles, the metacyc or uniprot databases, and lastly, we enriched each enzymatic step in said pathway with as much information as possible. Information that was added included for instance: GPR, reaction localization, EC numbers, a confidence score, possible cofactors and inhibitors and cross references to other databases such as KEGG, BIGG and MetaNetX. For each metabolite involved in these reactions, we defined the stoichiometry, charge and elemental formula, either based on the corresponding entries in the BiGG database or on clues from literature.

If reactions from a pathway were present in the draft, we checked their constraints and directionality. This was necessary as many of the irreversible reactions in the draft reconstruction seemed to have been ‘inverted’ when compared to the corresponding reactions in the reference databases, which made flux through them impossible in normal growth conditions.

The energy metabolism and methane oxidation were curated first. Except for the reaction representing the sMMO, all reactions were newly constructed, as they were absent in the draft. Then, in order to achieve sensible FBA solutions for growth on methane, the central carbon metabolism, amino acid and nucleotide biosynthesis pathways were manually curated. Simultaneous to the manual curation a metabolic pathway map was created in Escher, which helped us to maintain a visual checklist of curated pathways.

The automated draft contained a rudimentary, generic biomass reaction, which only accounted for the production of proteins, DNA and RNA, but not for the biosynthesis of a cell wall and cell membrane, the maintenance of a cofactor pool, the turnover of trace elements or the energetic costs associated with growth. After calculating a more specific biomass composition for *M. capsulatus*, further pathway curation was necessary to achieve growth *in silico*. This included the biosynthesis pathways of fatty acids (even, odd and cyclopropane varieties), phospholipids, coenzyme A, Ubiquinol, Lanosterol, FAD, FMN, Riboflavin, NAD and NADP, Heme, Glutathione, Cobalamin, Thiamine, Myo-Inositol, and Lipopolysaccharides.

To account for the reported differences in ammonia assimilation of *M. capsulatus* when grown in the presence of excess ammonia versus the growth on either atmospheric nitrogen or nitrate, we curated the nitrogen metabolism including the oxidation of ammonia to nitrite via hydroxylamine, the reduction of nitrate and nitrite, the glutamine synthetase/ glutamate synthase reactions and the alanine dehydrogenase. Reversible degradation reactions producing ammonia that would potentially bypass the characterized ammonia assimilation pathways were constrained to be irreversible accordingly.

After we had enriched the annotations already in the draft with annotations from the metabolic models iJO1366 [52], iRC1080 [76], iJN678 [77] and iHN637 [78], they were converted into a MIRIAM-compliant format. As a final step, we manually added transport reactions to reflect the uptake energetics of cofactors.

Throughout the reconstruction process, we iteratively tested and validated the reconstruction. For instance, we checked the mass and charge balance of each reaction, attempting to manually balance those that weren’t balanced. In the majority of cases metabolites were missing formula or charge definitions. In order to remove energy generating cycles, problematic reactions were manually constrained to be irreversible. Validation was carried out against growth data [19], which was also the point of reference for the parameter fitting.

### Biomass Composition

For the principal components of biomass, measurements were made available through the website of our collaborators Unibio [23]. This included specifically the content of RNA (6.7%), DNA (2.3%), crude fat (9.1%), and glucose (4.5%) as a percentage of the cell dry weight. We did not use the percentage values for crude protein (72.9%) and N-free extracts (7.6%) as these measurements are fairly inaccurate relying on very generalized factors. The percentage value of Ash 550 (8.6%) measurements was inconsistent with the sum of its individual components (4.6%) and was hence excluded.

On the homepage of UniBio, we were also able to find g/kg measurements of all amino acids except for glutamine and asparagine, trace elements and vitamins, which could directly be converted into mmol/g DW. However, we omitted some of data: The stoichiometries for Selenium, Cadmium, Arsenic, Lead and Mercury were not included in the biomass reaction as their values were negligibly small. Beta-Carotene (Vitamin A) and Gama-Tocopherol (Vitamin E) were omitted because no genes were found supporting their biosynthesis, in addition to both being reportedly below the detection limit [79].

For the lack of better measurements, and assuming that *M. buryatense* and *M. capsulatus* are more similar than *M. capsulatus* and *E. coli,* the stoichiometries of glutamine and asparagine, intracellular metabolites such as ribulose-5-phosphate, organic cofactors such as coenzyme A, and cell wall components such as LPS were introduced from de la Torre et al. [20].

Using the GC content calculated from the genome sequence [22] and the percentage measurements from Unibio for RNA and DNA, we were able to calculate the stoichiometries of all individual nucleobases.

Unibio’s measurements that 94% of the crude fat portion were fatty acids conflicted with previously published results, which indicated that in fact phospholipids are likely to be the main lipid component in *M. capsulatus* [80, 81]. Thus, we assumed 94% of the crude fat to be phospholipids. This meant that 6% of the crude fat was composed of fatty acids, the distributions of which were again provided by Unibio. However, in order to calculate the stoichiometry of each fatty acid we recalculated the distribution to exclude the unidentified part. Makula had also measured the composition of the phospholipid pool itself [80], from which we calculated the corresponding stoichiometries for phosphatidylethanolamine, phosphatidylcholine, phosphatidylglycerol and cardiolipin.

Bird and colleagues had reported the percentage of squalene and sterols of dry weight [82], which we converted into mmol/g DW without further corrections. Since the genes for hopanoid synthesis were identified we included diplopterol in the biomass reaction [22, 83]. For a lack of more detailed measurements we estimated a similar contribution to the overall composition as squalene. We specifically used lanosterol to represent the general class of sterols in the biomass reaction, since we had only been able to reconstruct the synthesis pathway of lanosterol and since lanosterol is a key precursor metabolite in the biosynthesis of all other sterols.

The growth associated maintenance energy requirements were calculated in accordance with protocol procedures [16].

### Transport Reactions

The identification of transport reactions involved the two databases for transport proteins, the Transporter Classification Database (TCDB) [84] and the Transport Database (TransportDB) [71], and two computational tools, PSORTb v3.0 [69] and BLASTp [70]. We employed the following semi-automatic way of inferring the putative function of transport proteins in *M. capsulatus*.

Using the protein sequences in AE017282.2_protein.faa [22], we determined the subcellular location of each protein using PSORTb v3.0. We filtered the results and focused only on proteins with a final score larger than 7.5, which the authors of PSORTb consider to be meaningful. We combined this list with the *M. capsulatus* specific entries from the TransportDB, which allowed us to use the PSORT-scores as an additional measure of confidence. At this point, 242 putative transport proteins were identified. From this list we then selected all proteins which were predicted to transport metabolites and were already included in the model, which reduced the number to 133. Since for many of the entries, the exact mechanism of transport is unknown, we combined the previously selected transporters with the results from running BLAST against the TCDB. The e-value and bitscore from BLAST provided yet another measure to confidently assess the correctness of the automatic TransportDB predictions, and the Transporter Classification-Numbers (TC-numbers) allowed us to gather information on the mechanism of transport. This led to a list of 97 transport proteins with high confidence, which was filtered once more as follows.

We checked the general description for a given specific TC-number, and then considered the BLAST result to read about a given transporter in more detail, especially with regards to finding the specific substrates. When we were able to identify the corresponding transport reaction in the BiGG database, we incorporated only the simplest, smallest set of reactions. In cases of conflicts between our own BLAST results and the automatic TransportDB transporter annotation, we preferentially trusted the BLAST results. Thus we ultimately added 75 transporter-encoding genes connected via GPR to 56 distinct transport reactions.

### Ratio Constraint

To identify which mode of electron transfer is active in *M. capsulatus*, we fit the solutions of FBA using iMcBath to measured values [19]. The authors had experimentally determined the O_2_/CH_4_ ratio and the growth yield of *M. capsulatus* in several conditions. They varied the nitrogen source using KNO_3_, NH_4_CL, and both simultaneously; the concentration of copper in the medium, which directly affects the activity of either sMMO or pMMO; and whether the culture was oxygen or carbon limited.

Since each electron transfer mode respectively is represented by the flux through one specific reaction in the model (PMMOipp, PMMODCipp, CYOR_q8ppi), we were able to investigate each simply by constraining the corresponding reaction.

Secondly, we accounted for the differential expression of ADH in the presence of excess NH_4_ in the medium versus the GS/GOGAT in the presence of NO_4_ or N_2_ by blocking the corresponding reactions accordingly [53].

Several studies have shown that NH_4_ is a co-metabolic substrate to the methane monooxygenases in *M. capsulatus* leading to the production of hydroxylamine first and nitrite later [54, 74, 85]. Hence, when simulating the growth on NH_4_ we assumed that varying ratios ***r*** of the nitrogen taken up would eventually be converted into nitrite (Figure 7).

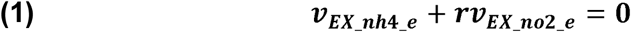

### Stoichiometric Modeling

The reactome of an organism can be represented mathematically as a stoichiometric matrix ***𝒮***. Each row of ***𝒮*** corresponds to a unique compound, while each column corresponds to a metabolic reaction. Hence the structure of the matrix for an organism with ***m*** compounds and n reactions equals ***m X n***. The values in each row denote the stoichiometric coefficients of that metabolite in all reactions. The coefficients are either negative for compounds that are educts, positive for those that are products, or zero for those that are not involved in a given metabolic reaction.

The vector ***v*** of length ***n*** contains as values the turnover rates of each reaction. These rates are commonly referred to as fluxes and are given the unit mmol gDW^-1^ h^-1^. Vector ***v*** is also referred to as the flux vector,

## Declarations

### Ethics approval and consent to participate

“Not applicable”

### Consent for publication

“Not applicable”

### Authors’ contributions

CL collected and analysed the data, reconstructed the metabolic model, drafted and revised the manuscript. LP and SBJ provided feedback on the metabolic behaviour of *M. capsulatus*, discussed improvements to the model and revised the manuscript. KVG, MJH and NS conceived and supervised the study and revised the manuscript critically. All authors read and approved the manuscript.

## Acknowledgements

The authors would like to acknowledge Budi Juliman Hidayat, Subir Kumar Nandi, John Villadsen for sharing their experiences with *M. capsulatus*; Mitch Pesesky, David Collins, Marina Kalyuzhnaya, Ilya Akberdin, and Sergey Stolyar for insightful discussions at the GRC C1 Conference 2016; and Kristian Jensen, Joao Cardoso, and Marta Matos for their help with software problems and feedback on the implementation.

